# Targeting interleukin-1 for reversing fat browning and muscle wasting in infantile nephropathic cystinosis

**DOI:** 10.1101/2021.02.05.429989

**Authors:** Wai W Cheung, Sheng Hao, Ronghao Zheng, Zhen Wang, Alex Gonzalez, Ping Zhou, Hal M Hoffman, Robert H Mak

## Abstract

**Background:** *Ctns*^−/−^ mice, a mouse model of infantile nephropathic cystinosis, exhibit hypermetabolism with adipose tissue browning and profound muscle wasting. Inflammatory cytokines such as IL-1 trigger inflammatory cascades and play an important role in the pathogenesis of cachexia. Anakinra is an FDA-approved IL-1 receptor antagonist that blocks IL-1 signaling and may provide targeted novel therapy.

**Methods:** *Ctns*^−/−^ mice were bred to *Il6* ^−/−^ and *Il1β* ^−/−^ mice. *Ctns*^−/−^ mice and wild type control were treated with anakinra (2.5 mg.kg.day, IP) or saline as vehicle for 6 weeks. We quantitated total fat mass and studied expression of molecules regulating adipose tissue browning. We measured gastrocnemius weight, total lean mass content, muscle function (grip strength and rotarod activity), muscle fiber size, muscle fatty infiltration and expression of molecules regulating muscle metabolism. We also evaluated the effects of anakinra on the muscle transcriptome.

**Results:** Il-1β deficiency or treatment with anakinra normalized food intake and weight gain, fat and lean mass content, metabolic rate and muscle function in *Ctns*^−/−^ mice. Anakinra also diminished molecular perturbations of energy homeostasis in adipose tissue and muscle, specifically, aberrant expression of beige adipose cell biomarkers (UCP-1, CD137, Tmem26 and Tbx1) and molecules implicated in adipocyte tissue browning (Cox2/Pgf2α, Tlr2, Myd88 and Traf6) in inguinal white adipose tissue in *Ctns*^−/−^ mice. Moreover, anakinra normalized gastrocnemius weight and fiber size as well as attenuated muscle fat infiltration in *Ctns*^−/−^ mice. This was accompanied by correction of the increased muscle wasting signaling pathways (increased protein content of ERK1/2, JNK, p38 MAPK and NF-κB p65 and gene expression of Atrogin-1 and Myostatin) and the decreased myogenesis process (decreased gene expression of MyoD and Myogenin) in gastrocnemius of *Ctns*^−/−^ mice. Finally, anakinra normalized or attenuated 12 of those top 20 differentially expressed muscle genes in *Ctns*^−/−^ mice.

**Conclusions:** Anakinra attenuates adipose tissue browning and muscle wasting in *Ctns*^−/−^ mice. IL-1 receptor blockade may represent a novel targeted treatment for cachexia in patients with infantile nephropathic cystinosis.

## INTRODUCTION

Cystinosis is a multisystem genetic disorder characterized by the accumulation of cystine in different tissues and organs. Infantile nephropathic cystinosis (INC) is the most common and severe form of cystinosis.^3^ Patients suffering from INC exhibit signs and symptoms of renal Fanconi syndrome and chronic kidney disease in early childhood.^4^ Metabolic abnormalities such as cachexia are common complications in patients with INC, which are associated with poor quality of life and mortality and for which there is no current therapy.^3,4^ Increased expression of inflammatory cytokines, including IL-1β, has been implicated in the etiology of cachexia and muscle wasting associated with different disease processes. Consistent with this finding, patients with cystinosis had higher concentrations of circulating IL-1β compared with controls.^6^ PBMCs from patients with cystinosis revealed a significant increase in IL-1β transcript levels compared with controls.^6^ Recent evidence in preclinical models suggests that blockade of IL-1 signaling may be a logical therapeutic target for chronic disease-associated muscle wasting since IL-1β activates NF-κβ signaling and induces expression of IL-6 and atrogin-1 in C2C12 myocytes.^7,8^ Anakinra is an IL-1 receptor antagonist which blocks both IL-1α and IL-1β.^10^ Duchene muscular dystrophy (DMD) is an X-linked muscle disease characterized by muscle inflammation that is associated with an increased circulating serum levels of IL-1β. Subcutaneous administration of anakinra normalized muscle function in a mouse model of DMD.^11^ Similarly, serum IL-1β is elevated in hemodialysis patients and a 4-week treatment with anakinra was shown to be safe in these patients while significantly reducing markers of systemic inflammation such as CRP and IL-6.^12^ In this study, we investigated *Ctns*^−/−^ mice as an established model of INC in which we have previously shown hypermetabolism with adipose tissue browning and profound muscle wasting by 12 months of age.^5^ We explored the role of inflammatory cytokines in *Ctns*^−/−^ mice, and evaluated the efficacy of IL-1 targeted therapy on adipose tissue browning and muscle wasting. Anakinra was FDA-approved for the treatment of rheumatoid arthritis in 2001 and is safe and an effective therapeutic option in a variety of diseases including diseases involving muscle. It may be repurposed as novel therapy for cachexia in INC.

## METHODS

### Study design

All animal work was conducted in compliance and approved by the Institutional Animal Care and Use Committee (IACUC) at the University of California, San Diego. Wild-type (WT), *Il6* ^−/−^ and *Il1β* ^−/−^, *Ctns^−/−^, Ctns^−/−^ Il6* ^−/−^ and *Ctns^−/−^Il1β* ^−/−^ mice were on the same c57BL/6 genetic background. *Ctns*^−/−^ mice were kindly provided by Professor Corinne Antignac and *Ctns^−/−^ Il6* ^−/−^ and *Ctns^−/−^Il1β* ^−/−^mice were generated by crossing *Ctns*^−/−^ mice with *Il6* ^−/−^ and *Il1β* ^−/−^ mice, respectively. Twelve-month old male mice were used for all of the following 5 studies. Study 1 – We evaluated the gastrocnemius muscle IL-1*β* and IL-6 mRNA and protein content in *Ctns*^−/−^ mice and WT mice. Results were presented in Figure 1, A to D. Study 2 – We evaluated the effects of genetic deletion of IL-1*β* and IL-6 in *Ctns*^−/−^ mice. We compared *ad libitum* food intake and weight change in WT, *Ctns^−/−^, Ctns^−/−^ Il6* ^−/−^ and *Ctns^−/−^Il1β* ^−/−^ mice. The study period was 6 weeks. Results were shown in Figure 1, E & F. Study 3 – We evaluated the beneficial effects of genetic deletion of IL-1*β* and IL-6 in *Ctns*^−/−^ mice beyond nutritional effects by employing a pair-feeding strategy. The study period was 6 weeks. *Ctns*^−/−^ mice were fed *ad libitum* and then WT, *Ctns^−/−^ Il6* ^−/−^ and *Ctns^−/−^Il1β* ^−/−^ mice were fed the same amount of rodent diet based on the recorded food intake of *Ctns*^−/−^ mice. Results were shown in Figure 1, G to N. Study 4 - We evaluated the effects of anakinra in *Ctns*^−/−^ mice. WT and *Ctns*^−/−^ mice were given anakinra (2.5 mg/kg.day, IP) or vehicle (normal saline), respectively. The study period was 6 weeks. All mice were fed *ad libitum*. We compared food intake and weight change in all groups of mice. Results were shown in Figure 2, A & B. Study 5 - We evaluated the metabolic effects of anakinra in *Ctns*^−/−^ mice beyond nutritional stimulation by employing a pair-feeding strategy. WT and *Ctns*^−/−^ mice were given anakinra (2.5 mg/kg.day, IP) or vehicle (normal saline), respectively. The study period was 6 weeks. Vehicle-treated *Ctns*^−/−^ mice were fed *ad libitum* while all other groups of mice were fed the same amount of rodent diet based on the recorded food intake of vehicle-treated *Ctns*^−/−^ mice. Results were shown in Figure 2, E to G & Figures, 3 to 6 as well as Supplemental Figure 1 & 2.

**Figure 1:**
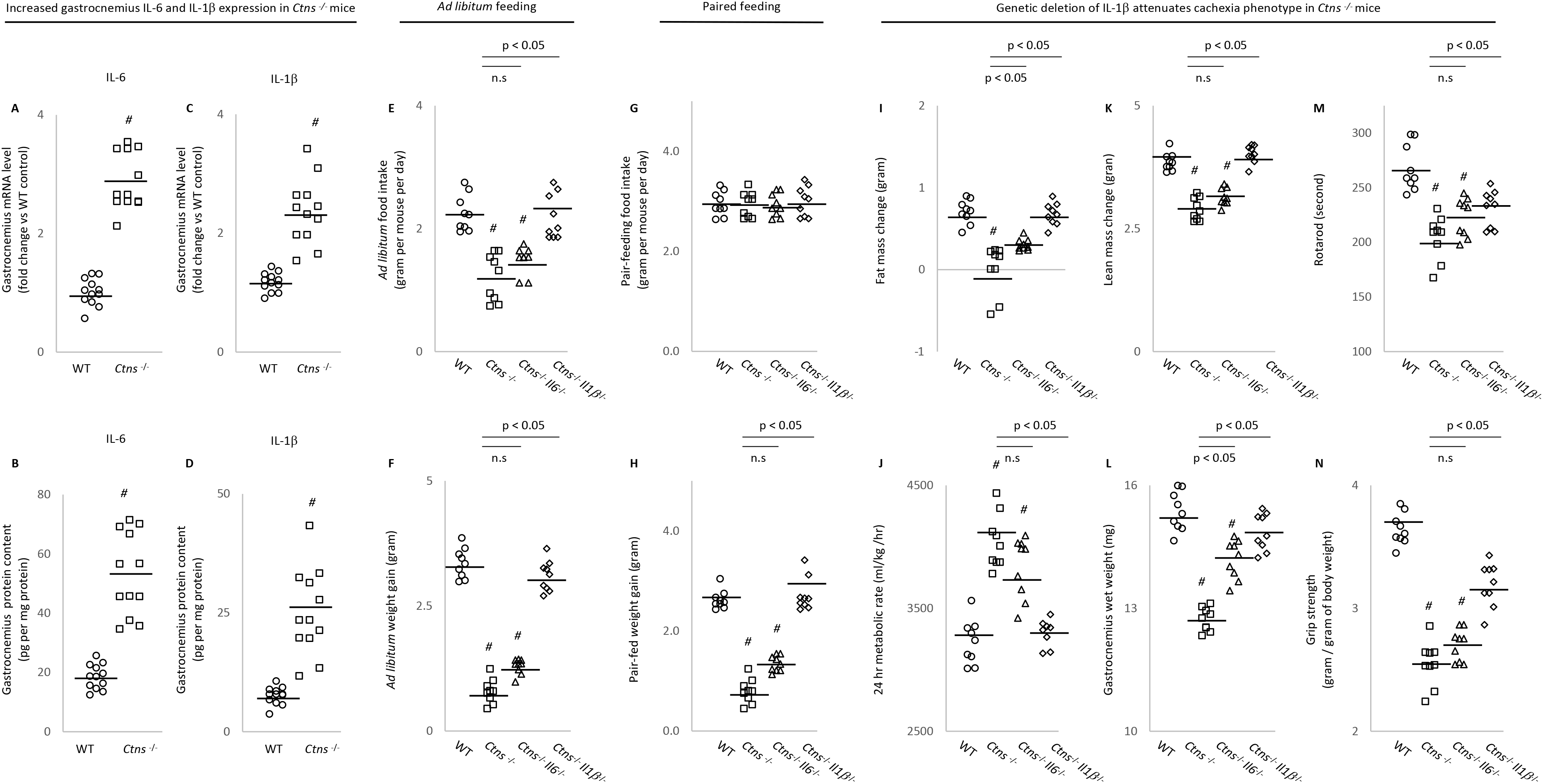
Increased muscle mRNA and protein content of IL-6 and IL-1β in *Ctns*^−/−^ mice and genetic depletion of IL-1β provides a better rescue of cachexia in *Ctns*^−/−^ mice compared to IL-6 deficiency. Results of 3 different experiments were shown. For the 1^st^ study, we compared gene expression and protein content of IL-6 and IL-1β in gastrocnemius muscle in 12-month old *Ctns*^−/−^ and WT mice. Data were expressed as mean ± SEM. Results of *Ctns*^−/−^ mice were compared to WT mice (A to D). For the 2^nd^ study, we compared the metabolic effects of genetic deletion of IL-6 and IL-1*β* in *Ctns*^−/−^ mice. Four groups of mice were included, *i.e., Ctns*^−/−^, *Ctns*^−/−^*Il6*^−/−^ and *Ctns^−/−^ Il1β* ^−/−^ as well as WT mice. All mice were 12-month old and were fed *ad libitum* and the study period was 6 weeks. Food intake and weight gain in mice were compared (E and F). For the 3^rd^ study, we investigated the metabolic effects of genetic deletion of IL-6 and IL-1*β* in *Ctns*^−/−^ mice beyond nutritional stimulation by employing a pair-feeding approach. Specifically, *Ctns*^−/−^ mice were fed *ad libitum* for 6 weeks while *Ctns^−/−^Il6^−/−^, Ctns^−/−^ Il1β* ^−/−^ and WT mice were pair-fed to that of *Ctns*^−/−^ mice (G). Weight gain, fat and lean content, 24-hr oxygen consumption, left gastrocnemius wet weight and *in vivo* muscle function (rotarod and grip strength) was measured in mice (H to N). Results of 2^nd^ and 3^rd^ study were expressed as mean ± SEM. Results of *Ctns*^−/−^, *Ctns^−/−^ Il6*^−/−^ and *Ctns^−/−^ Il1β* ^−/−^ were compared to WT mice significance represented by *#)*. In addition, results of *Ctns^−/−^ Il6*^−/−^ and *Ctns^−/−^ Il1β* ^−/−^ mice were also compared to *Ctns*^−/−^ mice, respectively (E to N) and statistical significance (p value) is shown above bar.

**Figure 2:**
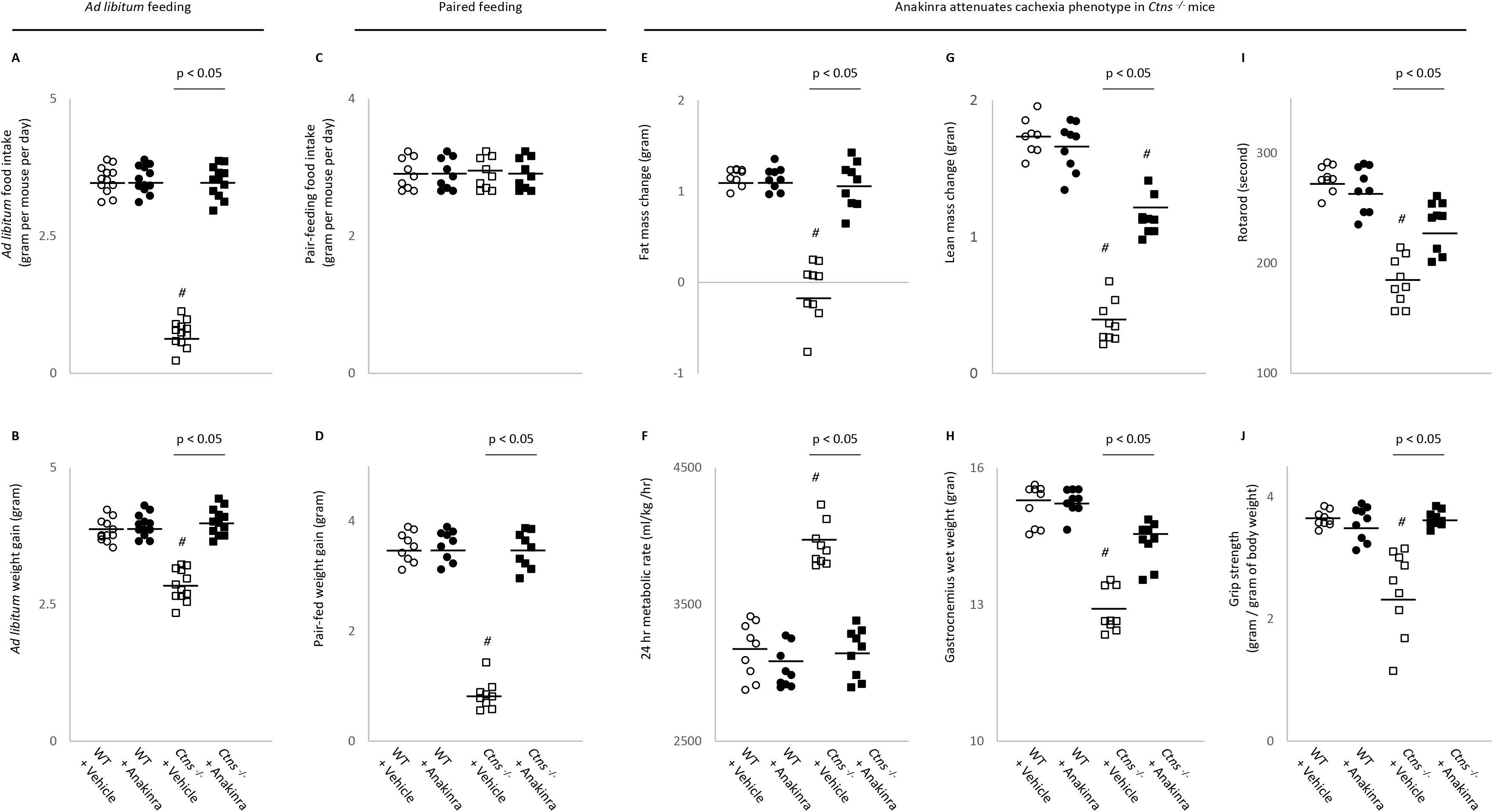
Anakinra attenuates cachexia in *Ctns*^−/−^ mice. Twelve-month old WT and *Ctns*^−/−^ mice were treated with anakinra (2.5 mg.kg.day, IP) or normal saline as a vehicle control for 6 weeks. All mice were fed *ad libitum* for 6 weeks and food intake as well as weight were recorded (A & B). In another experiment, to assess the beneficial effects of anakinra beyond its nutritional effects, we employed a pair-feeding strategy. Vehicle-treated *Ctns*^−/−^ mice were given an *ad libitum* amount of food whereas other groups of mice were given an equivalent amount of food (C). Weight gain, fat and lean content, 24-hr oxygen consumption, left gastrocnemius wet weight and *in vivo* muscle function (rotarod and grip strength) was measured in the mice (D to J). Data are expressed as mean ± SEM. Results of vehicle-treated *Ctns*^−/−^ mice were compared to vehicle-treated WT mice while results of anakinra-treated *Ctns*^−/−^ mice were compared to that of anakinra-treated WT mice (^*#*^ p < 0.05). In addition, results of anakinra-treated *Ctns*^−/−^ mice were compared to vehicle-treated *Ctns*^−/−^ mice and specific *p* value is shown above bar.

### Serum and blood chemistry

BUN and serum concentration of bicarbonate was measured (**Supplemental Table 1S**). Serum creatinine were analyzed by LC-MS/MS method.^13^

### Body composition, metabolic rate and *in vivo* muscle function

Body composition (for lean and fat content) of mice were measured by quantitative magnetic resonance analysis (EchoMRI-100™, Echo Medical System) twice, prior to the initiation of the study as well as at the end of the study and change of total body fat and lean mass in individual mouse was calculated. Twenty-four-hour metabolic rate (VO2) of mice were measured using Oxymax indirect calorimetry (Columbus Instrument) at the end of the study. At the end of the study, muscle function (grip strength and rotarod activity) in mice was assessed using a grip strength meter (Model 47106, UGO Basile) and rotarod performance tool (model RRF/SP, Accuscan Instrument), respectively.^5,14^

### Protein assay for muscle and adipose tissue

Mice were sacrificed at the end of the study and gastrocnemius muscle, inguinal white adipose tissue (WAT) and intercapsular brown adipose tissue (BAT) were dissected and processed in a homogenizer tube (USA Scientific, catalog 1420-9600) containing ceramic beads (Omni International, catalog 19-646) using a Bead Mill Homogenizer (Omni International). Protein concentration of tissue homogenate was assayed using Pierce BAC Protein Assay Kit (Thermo Scientific, catalog 23227). Uncoupling protein (UCP) protein content as well as adenosine triphosphate (ATP) concentration in adipose tissue and muscle homogenates were assayed. Protein concentration of phospho-ERK 1/2 (Thr202/Ty2r204) and total ERK 1/2, phospho-JNK (The183/Tyr185) and Total JNK, phospho-p38 MAPK (Thr180/Tyr182) and total p38 MAPK, NF-κB p65 (phospho-Ser536) and total NF-κB p65 as well as IL-6 and IL-1β in muscle homogenates were assayed (**Supplemental Table 1S**).

### Gastrocnemius wet weight, fiber size and fatty infiltration

Left gastrocnemius muscle of mice was dissected at the end of the study. Wet weight of left gastrocnemius muscle was recorded. Subsequently, we measured muscle fiber cross-sectional area of gastrocnemius muscle. using ImageJ software (https://rsbweb.nih.gob/ij/).^5,14^ In addition, portion of dissected right gastrocnemius muscle samples were incubated with Oil Red O (Oil Red O Solution, catalog number O1391-250 ml, Sigma Aldrich). Detailed procedures for Oil Red O staining were in accordance with published protocol.^15^ We followed a recently established protocol to quantify muscle fat infiltration. Acquisition and quantification of images were analyzed using ImageJ software.^16^

### Muscle RNAseq analysis

Previously, we performed RNAseq analysis on gastrocnemius muscle mRNA in 12-month old *Ctns*^−/−^ mice versus age-appropriate WT mice.^14^ Detailed procedures for mRNA extraction, purification and subsequent construction of cDNA libraries as well as analysis of gene expression was published. We then performed Ingenuity Pathway Analysis enrichment tests for those differentially expressed muscle genes in *Ctns*^−/−^ mice versus WT mice, focusing on pathways related to energy metabolism, skeletal and muscle system development and function, and organismal injury and abnormalities. We identified the top 20 differentially expressed muscle genes in *Ctns*^−/−^ versus WT mice.^14^ In this study, we performed qPCR analysis for those top 20 differentially expressed gastrocnemius muscle genes in the different experimental groups.

### Quantitative real-time PCR

Total RNA from adipose and gastrocnemius muscle samples were isolated using TriZol (Life Technology) and reverse-transcribed with SuperScript III Reverse Transcriptase (Invitrogen). Quantitative real-time RT-PCR of target genes were performed using KAPA SYBR FAST qPCR kit (KAPA Biosystems). Expression levels were calculated according to the relative 2^−ΔΔCt^ method.^14^ All primers are listed (**Supplemental Table 2S**).

### Statistical analysis

Continuous variables are expressed as mean ± S.E.M. We assessed the statistical significance of differences between groups using two-sample t-tests. All tests were two-sided. A *p* value less than 0.05 was considered significant. Statistical analyses were performed using SPSS software version 16.0 for Macintosh. *#* represents a statistically significant (*p* < 0.05) difference from WT control mice. Bars with specific *p* values between groups represents a statistical significance from *Ctns*^−/−^ control mice.

## RESULTS

### Increased muscle IL-6 and IL-1β mRNA and protein content in *Ctns*^−/−^ mice

We chose to study *Ctns*^−/−^ mice at 12 months of age based on our previous data demonstrating a significant cachexia phenotype. As expected, all of the mice were uremic and had elevated creatine, but normal bicarbonate levels serum (**Supplemental Table 3S**). In order to investigate the role of inflammatory cytokines in cachexia and muscle wasting, we measured gastrocnemius muscle mRNA and protein content of IL-6 and IL-1β in 12-month old *Ctns*^−/−^ mice and age-matched WT controls. Gastrocnemius mRNA and protein content of both IL-6 and IL-1β were significantly elevated in *Ctns*^−/−^ mice relative to WT mice (**Figure 1, A-D**).

### Genetic deletion of IL-1β normalizes cachexia in *Ctns*^−/−^ mice

We then employed a genetic approach to determine whether IL-6 and IL-1β deficiency would affect the cachexia phenotype in *Ctns*^−/−^ mice. Twelve-month old *Ctns*^−/−^, *Ctns*^−/−^*Il6*^−/−^ and *Ctns^−/−^Il1β* ^−/−^ mice were all uremic and had elevated creatinine levels but normal bicarbonate levels (**Supplemental Table 4S**). All mice were fed *ad libitum* for 6 weeks and average daily food intake as well as final weight gain of the mice were recorded. We showed that genetic deletion of *Il1β* completely corrected anorexia and normalized weight gain in *Ctns*^−/−^ mice whereas deletion of *Il6* only partially rescued the phenotype in *Ctns*^−/−^ mice (**Figure 1, E&F**). Furthermore, to investigate the potential metabolic effects of genetic deficiency of *Il1β* and *Il6* beyond its nutritional effects, we employed a pair-feeding strategy so *Ctns*^−/−^ mice were fed *ad libitum* and then all other groups of mice were fed the same amount of rodent diet based on the recorded food intake of the *Ctns*^−/−^ group (**Figure 1G**). Despite receiving the same amount of total calorie intake as other groups of mice, cardinal features of cachexia phenotype such as decreased weight gain, decreased fat mass content and hypermetabolism (manifested as elevated oxygen consumption), decreased lean mass content and gastrocnemius weight as well as reduced muscle function (decreased rotarod and grip strength) were still prominent in *Ctns*^−/−^ mice (**Figure 1, H to N**). Importantly, parameters of cachexia phenotype were normalized relative to WT mice or significantly improved in *Ctns^−/−^Il1β* ^−/−^ mice compared to *Ctns*^−/−^ mice. However, *Ctns^−/−^Il6* ^−/−^ mice had only minimal non-significant improvement relative to *Ctns*^−/−^ mice.

### Anakinra attenuates cachexia and muscle wasting in *Ctns*^−/−^ mice

In order to better model a real-world approach in patients, we used a pharmacological approach to test the effects of IL-1β blockade in pre-established INC-associated cachexia and muscle wasting. Anakinra, an IL-1 receptor antagonist that blocks both IL-1α and IL-1β function was used to treat 12-month old *Ctns*^−/−^ mice and WT controls for 6 weeks compared to vehicle. All mice were fed *ad libitum*. Food intake and weight gain was normalized in anakinra treated *Ctns*^−/−^ mice (**Figure 2, A to B**). We also investigated the beneficial effects of anakinra in *Ctns*^−/−^ mice beyond appetite stimulation and its consequent body weight gain through the utilization of a pair-feeding approach. Daily *ad libitum* food intake for *Ctns*^−/−^ mice treated with vehicle was recorded. Following that, anakinra treated *Ctns*^−/−^ mice as well as WT mice treated with anakinra or vehicle were food restricted such that the mice ate an equivalent amount of food as vehicle treated *Ctns*^−/−^ mice (**Figure 2C**). Serum and blood chemistry of mice were measured (**Supplemental Table 5S**). Anakinra normalized uremia, serum creatine, weight gain, and metabolic rate, and significantly improved or normalized fat and lean mass content, gastrocnemius weight, as well as muscle function (grip strength and rotarod activity) in *Ctns*^−/−^ mice (**Figure 2, D to J**). Results of anakinra-treated *Ctns*^−/−^ mice were also compared to vehicle-treated *Ctns*^−/−^ mice. Anakinra normalized parameters of cachexia phenotype in *Ctns*^−/−^ mice compared to vehicle-treated *Ctns*^−/−^ mice.

### Anakinra normalizes energy homeostasis in adipose tissue and skeletal in *Ctns*^−/−^ mice

Adipose tissue (inguinal WAT and intercapsular BAT) and gastrocnemius protein content of UCPs was significantly increased in *Ctns*^−/−^ mice compared to WT mice (**Figure 3, A, C & E**). In contrast, ATP content in adipose tissue and skeletal muscle was significantly decreased in *Ctns*^−/−^ mice compared to WT mice (**Figure 3, B, D & F**). Anakinra normalized UCPs and ATP content of adipose tissue and muscle in *Ctns*^−/−^ mice.

**Figure 3:**
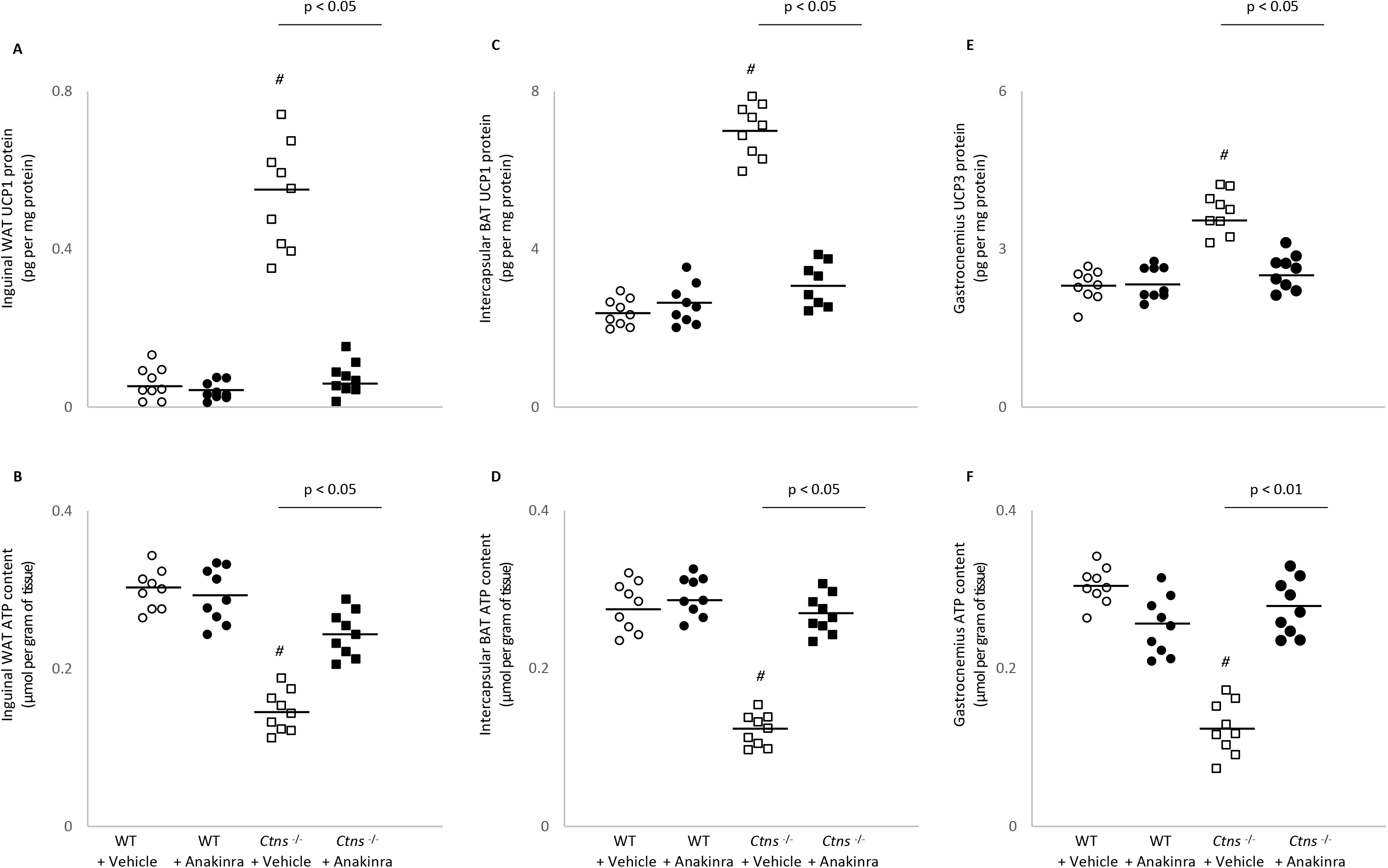
Anakinra ameliorates energy homeostasis in adipose tissue and skeletal in *Ctns*^−/−^ mice. UCPs and ATP content in adipose tissue (inguinal white adipose tissue and brown adipose tissue) and gastrocnemius muscle were measured. Results are analyzed and expressed as in Figure 2.

### Anakinra ameliorates browning of adipose tissue in *Ctns*^−/−^ mice

Beige adipose cell surface markers (CD137, Tmem26 and Tbx1) mRNA levels were elevated in inguinal WAT in *Ctns*^−/−^ mice relative to WT mice (**Figure 4, A to C)**. This was accompanied by an increased UCP-1 protein content in inguinal WAT shown in **Figure 3A**, which is a biomarker for beige adipocytes, and usually undetectable in WAT. Anakinra normalized the elevated protein content of inguinal WAT UCP-1 as well as increased mRNA expression of CD137, Tmen and Tbx1 in *Ctns*^−/−^ mice (**Figure 3A and Figure 4, A to C**). Anakinra also ameliorated expression of important signature molecules implicated in WAT. Activation of the Cox2/Pgf2α signaling pathway and chronic inflammation stimulate biogenesis of WAT browning. Significantly increased inguinal WAT mRNA expression of Cox2 and Pgf2α was found in *Ctns*^−/−^ mice compared to WT mice (**Figure 4, D and E**). Moreover, inguinal WAT levels of toll-like receptor 2 (Tlr2) as well as myeloid differentiation primary response 88 (MyD88) and tumor necrosis factor receptor-associated factor 6 (Traf6) were upregulated in *Ctns*^−/−^ mice compared to WT mice (**Figure 4, F to H**) and anakinra significantly reduced inguinal WAT gene expression of Cox2/Pgf2α as well as genes in the toll-like receptor pathway in *Ctns*^−/−^ mice.

**Figure 4:**
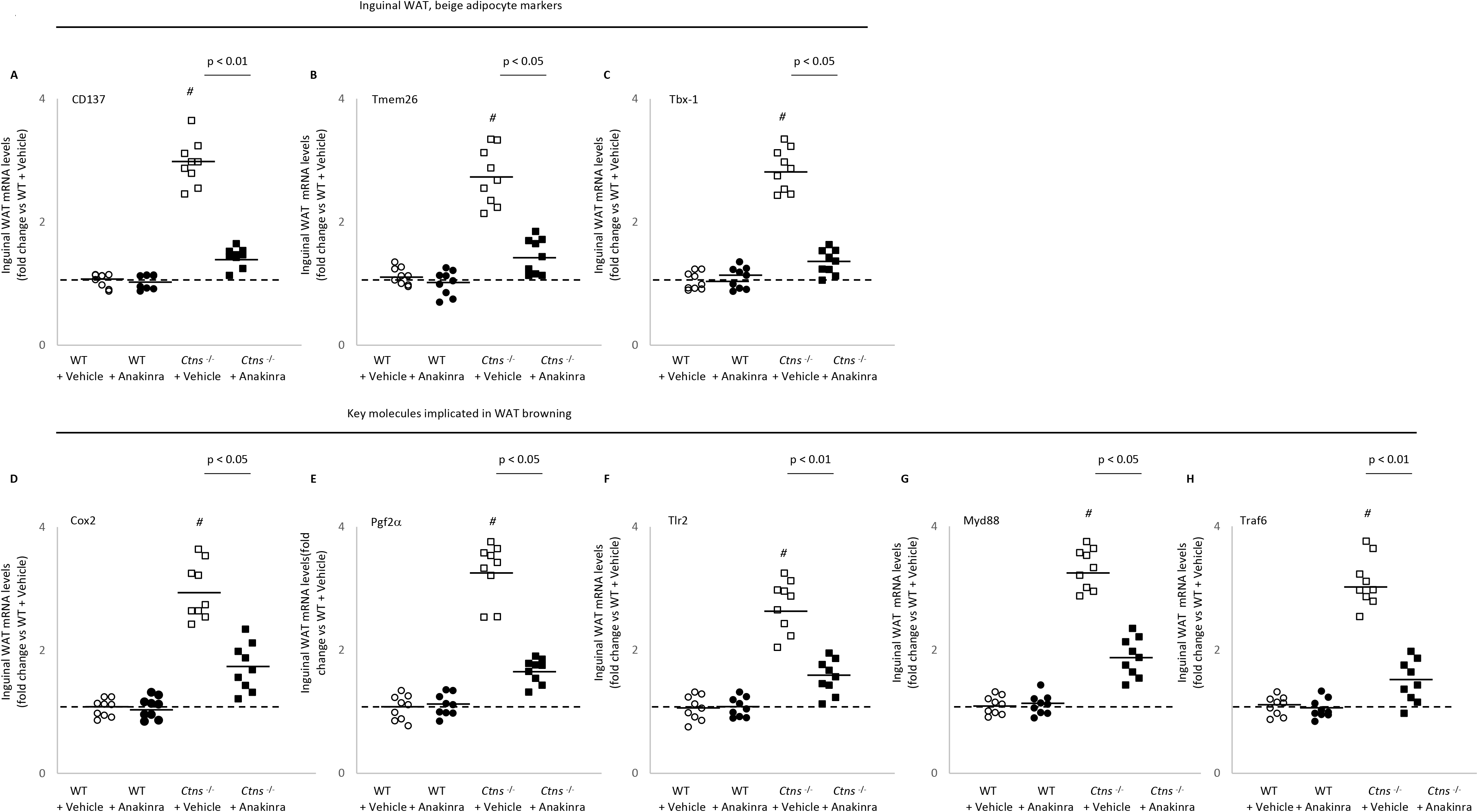
Anakinra attenuates adipose tissue browning in *Ctns*^−/−^ mice. Gene expression of beige adipocyte markers (CD137, Tmem26 and Tbx-1) in inguinal white adipose tissue was measured. Gene expression of Cox2 signaling pathway (Cox2 and Pgf2α) and toll like receptor pathway (Tlr2 and MyD88) in inguinal white adipose tissue was also measured. Gene expression was measured by qPCR. Final results were expressed in arbitrary units, with one unit being the mean level in vehicle-treated WT mice. Results are analyzed and expressed as in Figure 2.

### Anakinra improves muscle fiber size and attenuates muscle fat infiltration in *Ctns*^−/−^ mice

We evaluated the effect of anakinra on skeletal muscle morphology in *Ctns*^−/−^ mice. Representative results of muscle sections are illustrated (**Figure 5, A to D**). Anakinra significantly improved average cross-sectional area of gastrocnemius fibers in *Ctns*^−/−^ mice (**Figure 5E**). We also evaluated fat infiltration of the gastrocnemius muscle in *Ctns*^−/−^ mice. Representative results of Oil Red O staining of muscle section are shown (**Figure 5, F to I**). We quantified intensity of muscle fat infiltration in mice and observed that anakinra attenuated muscle fat infiltration in *Ctns*^−/−^ mice (**Figure 5J**).

**Figure 5:**
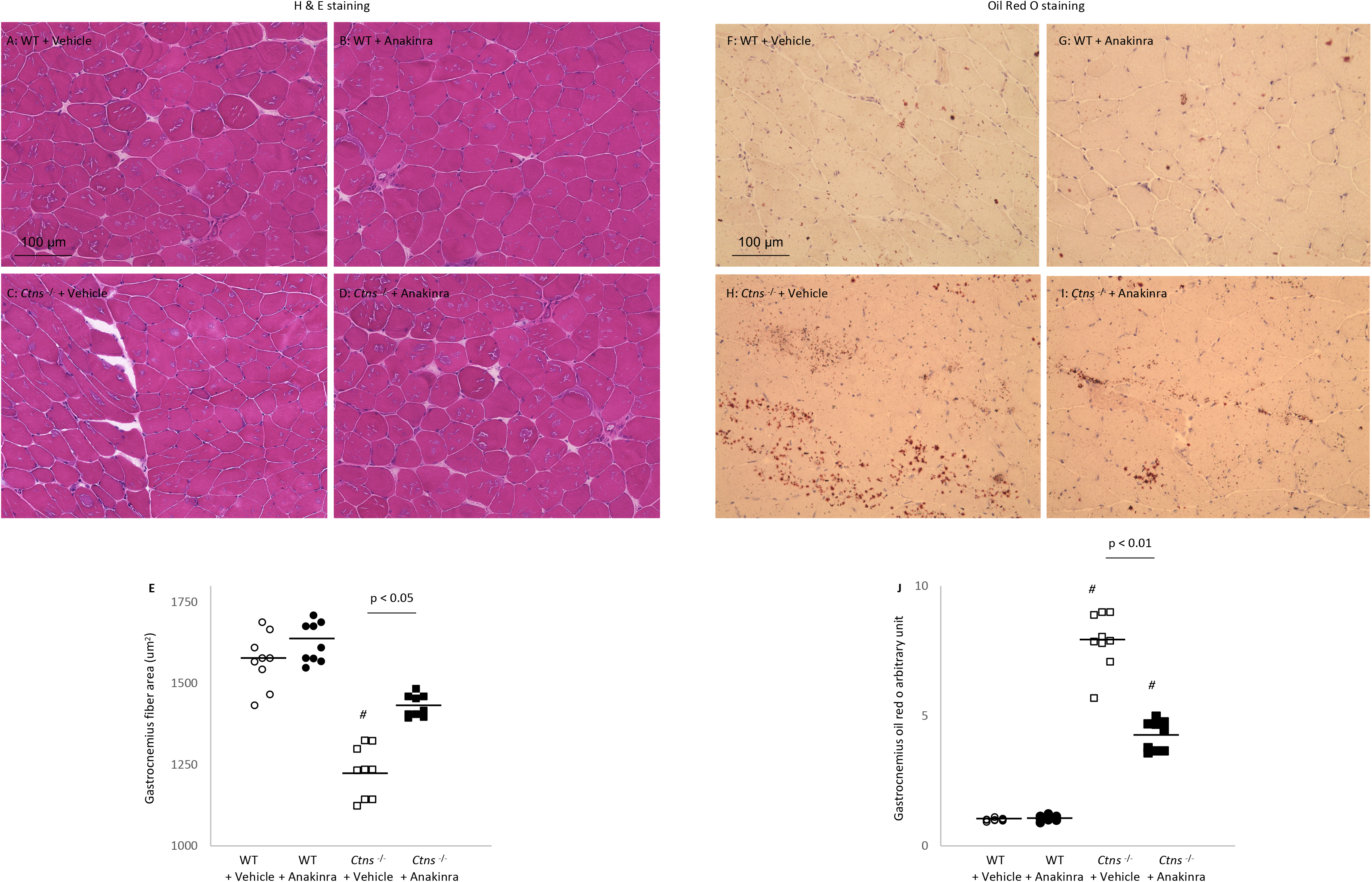
Anakinra normalizes muscle fiber size and attenuates muscle fat infiltration in *Ctns*^−/−^ mice. Representative photomicrographs of gastrocnemius with H&E staining (A-D). Average gastrocnemius cross-sectional area was measured (E). Visualization of quantification of fatty infiltration by Oil Red O analysis in gastrocnemius muscle (F-J). Final results were expressed in arbitrary units, with one unit being the mean staining intensity in vehicle-treated WT mice. Difference among various groups of mice were analyzed as in Figure 2.

### Anakinra attenuates signaling pathways implicated in muscle wasting in *Ctns*^−/−^ mice

Muscle relative phospho-ERK 1/2 (Thr202/Tyr204), phospho-JNK (Thr183/Tyr185) and phospho-p38 MAPK protein content (Thr180/Tyr182) was significantly increased in *Ctns*^−/−^ mice compared to WT mice (**Figure 6, A-C**). In addition, the ratio of muscle phospho-NF-κB p65 (phosphorylated S536) relative to total NF-κB p65 protein content was elevated in *Ctns*^−/−^ mice relative to WT mice (**Figure 6, D**). Importantly, anakinra attenuated or normalized phosphorylation of muscle ERK 1/2, JNK, p38 MAPK and NF-κB p65 protein content in *Ctns*^−/−^ mice. Administration of anakinra improved muscle regeneration and myogenesis by decreasing mRNA expression of negative regulators of skeletal muscle mass (Atrogin-1 and Myostatin) (**Figure 6, E and F**) as well as increasing muscle mRNA expression of pro-myogenic factors (MyoD and Myogenin) in *Ctns*^−/−^ mice (**Figure 6, G and H**).

**Figure 6:**
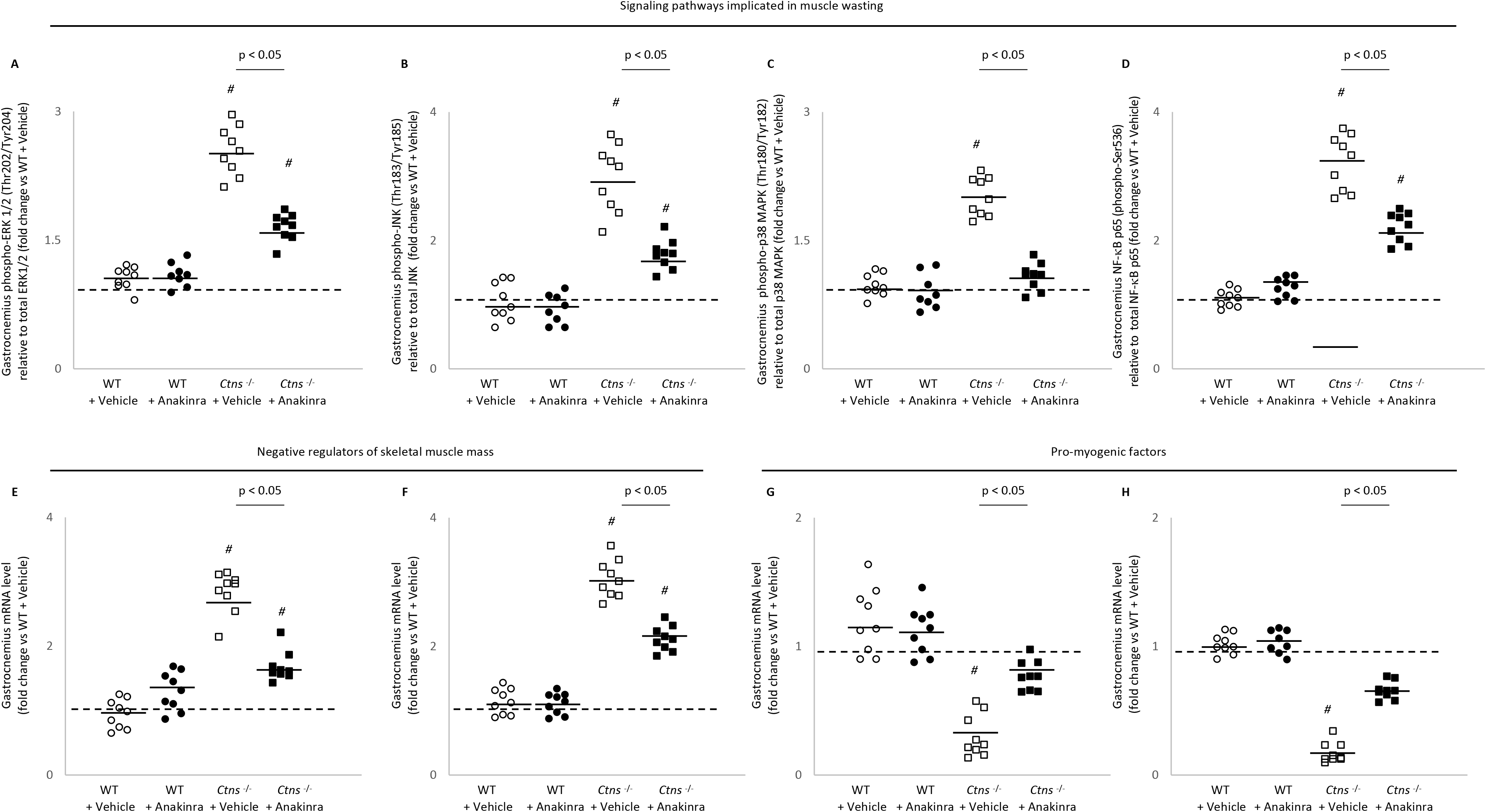
Anakinra attenuates signaling pathways implicated in muscle wasting in *Ctns*^−/−^ mice. Gastrocnemius muscle relative phospho-Akt (pS473) / total Akt ratio, relative phospho-ERK 1/2 (Thr202/Tyr204) / total ERK 1/2 ratio, relative phospho-JNK (Thr183/Tyr185) / total JNK ratio, relative phospho-p38 MAPK (Thr180/Tyr182) / total p38 MAPK and relative phosphorylated NF-κB p65 (Ser536) / total p65 ratio as well as relative phosphor Iκκα (Ser536) / total Iκκα ratio in mice. In addition, gastrocnemius muscle expression of interested genes was measured by qPCR. Final results were expressed in arbitrary units, with one unit being the mean level in vehicle-treated WT mice. Results are analyzed and expressed as in Figure 2.

### Molecular mechanism by RNAseq analysis

Recently, we profiled differential expression of gastrocnemius mRNA between 12-month old *Ctns*^−/−^ mice and WT mice using RNAseq analysis.^14^ Ingenuity Pathway Analysis enrichment tests identified the top 20 differentially expressed muscle genes in *Ctns*^−/−^ mice versus WT mice. The top 15 up-regulated genes were Ankdr2, Csrp3, Cyfip2, Fhl1, Ly6a, Mup1, Myl2, Myl3, Nlrc3, Sell, Sln, Spp1, Tnnc1, Tnni1, and Tpm3 whereas the top 5 down-regulated genes were Atf3, Cidea, Fos, Sncg and Tbc1d1 in *Ctns*^−/−^ mice relative to WT mice. We performed qPCR analysis for those top 20 differentially expressed muscle genes in the present study. Importantly, anakinra significantly reduced (Ankdr2, Csrp3, Cyfip2, Fhl1, Ly6a, Nlrc3, Tnnc1, and Tpm3) and significantly increased (Atf3, Cidea, Sncg, and Tbc1d1) muscle gene expression in *Ctns*^−/−^ mice relative to WT mice **(Supplemental Figure 1 & 2)**. Functional significance of each of these 12 genes is listed (**Table 1**). Nonsignificant changes were observed in Mup1, Myl2, Myl3, Sell, Sln, Spp1, Tnni1 and Fos.

**Table 1:** Anakinra normalizes or attenuated expression of important muscle genes that have been implicated in muscle wasting in *Ctns*^−/−^ mice. Previously, we studied differential expression of gastrocnemius mRNA between *Ctns*^−/−^ mice and WT mice using RNAseq analysis.^14^ We focus on pathways related to energy metabolism, skeletal and muscular system development and function, nervous system development and function as well as organismal injury and abnormalities. We performed qPCR analysis for top 20 differentially expressed muscle genes in the present study. Importantly, anakinra normalized (Ankdr2, Csrp3, Ly6a, Nlrc3, Tnnc1 and Tpm3, Atf3, Cidea, Sncg and Tbc1d1) and attenuated (Cyfip2 and Fhl1) muscle gene expression in *Ctns*^−/−^ mice relative to WT mice. Functional significance of each of these differentially expressed muscle genes is listed. DEG, differential expressed genes.

| **Upregulated DEG** | **Functional significance & reference** |
| --- | --- |
| Ankrd2 | implicated in mechanical stretch of skeletal muscle <sup>55,56</sup> |
| Csrp3 | associated with skeletal muscle dystrophy <sup>48</sup> |
| Cyfip2 | associated with muscle wasting <sup>49</sup> |
| Fhl1 | activates myostatin signaling and promotes atrophy in skeletal muscle <sup>50</sup> |
| Ly6a | associated with remodeling of the extracellular matrix during skeletal muscle regeneration <sup>54</sup> |
| Nlrc3 | implicated in skeletal muscle wasting by inhibiting cell proliferation and promoting cell apoptosis <sup>51</sup> |
| Tnnc1 | regulates straited muscle contraction <sup>52</sup> |
|  | implicated in cardiomyopathy pathogenesis and age-related skeletal muscle wasting <sup>52</sup> |
| Tpm3 | promotes slow myofiber hypotrophy and associated with generalized muscle weakness <sup>53</sup> |
| **Downregulated DEG** | **Functional significance** |
| Atf3 | impairs motor neuron survival and muscle innervation <sup>57</sup> |
| Cidea | Increases metabolic rates, lipolysis in brown adipose tissue and higher core temperature <sup>58</sup> |
| Sncg | Increases energy expenditure, particularly in brown and white adipose tissue <sup>59</sup> |
| | associated with cancer cachexia through the TGF-β signaling pathway <sup>59</sup> |
| Tbc1d1 | Impairs glucose transport in skeletal muscle <sup>60</sup> |
|  | associated with follistatin-induced muscle hypertrophy <sup>60</sup> |

## DISCUSSION

Previously, we described cachexia characterized by adipose tissue browning and muscle wasting in *Ctns*^−/−^ mice, an established mouse model of INC, but the etiology was not clear. Increased expression of inflammatory cytokines such as IL-6 and IL-1 have been implicated in the etiology of cachexia and muscle. Our studies using specific cytokine deficient mice and IL-1 targeted therapy suggest that IL-1 is a critical cytokine in INC-associated cachexia. The novel findings of this study are (i) Anakinra corrected anorexia, normalized weight gain, and attenuated hypermetabolism. (ii) Anakinra attenuated the expression of markers of adipose tissue browning as well as perturbed energy expenditure and thermogenesis. (iii) Anakinra attenuated muscle wasting, improved muscle function and corrected perturbations of aberrant expression of molecules implicated in muscle wasting. Together, our results suggest that anakinra may be an effective targeted treatment approach for cachexia in patients with INC.

IL-1β suppresses food intake, activates energy metabolism and reduces weight gain in experimental animals.^9,17^ IL-1β signals through appetite-regulating neuropeptides, such as leptin resulting in appetite suppression.^18^ We have demonstrated that elevated circulating level of leptin through the activation of melanocortin receptor 4 induces CKD-associated cachexia.^19^ For this present study, we showed that skeletal muscle IL-1β level was significantly increased in *Ctns*^−/−^ mice, and *Ctns^−/−^Il1β* ^−/−^ mice had normalized parameters of cachexia phenotype relative to control mice (**Figure 1**). Furthermore, we showed that anakinra improved anorexia and normalized weight gain in *Ctns*^−/−^ mice relative to control mice (**Figure 2**). Our results further highlight the beneficial effects of anakinra beyond food stimulation and accompanied weight gain. In pair-fed studies in which anakinra-treated *Ctns*^−/−^ mice and anakinra-treated WT mice were fed the same amount of food, cachexia was attenuated in anakinra-treated *Ctns*^−/−^ mice relative to control mice (**Figure 2**).

The basal metabolic rate accounts for up to 80% of the daily calorie expenditure by individual.^20^ Skeletal muscle metabolism is a major determinant of resting energy expenditure.^21,22^ IL-1β increases basal metabolic rate (as represented by an increase in resting oxygen consumption) in a dose-dependent manner.^23^ In our study, anakinra normalized the increased 24-hr metabolic rate in *Ctns*^−/−^ mice (**Figure 2J**). Adipose tissue UCP1 expression is essential for adaptive adrenergic non-shivering thermogenesis and muscle UCP3 level controls body metabolism.^24^ The energy generated when dissipating the proton gradient via upregulation of UCPs is not used for cellular ATP production or other biochemical processes but instead to produce heat.^25,26^ We showed that anakinra normalized muscle and adipose tissue UCPs and ATP content in *Ctns*^−/−^ mice (**Figure 3**). Blockade of IL-1 receptor signaling may also mitigate the metabolic dysfunction through leptin signaling. Infusion of leptin increased UCPs expression in skeletal muscle and adipose tissue.^27,28^

Perhaps our most novel finding is that anakinra reduced adipose tissue browning in *Ctns*^−/−^ mice. New evidence suggests a maladaptive role of adipose tissue browning in the context of cachexia.^29^ Activation of adipose tissue browning is associated with profound energy expenditure and weight loss in cachectic patients. Browning of adipose tissue increases energy expenditure and takes place before skeletal muscle atrophy in murine models of cancer and CKD-associated muscle wasting.^30,31^ We previously demonstrated adipose tissue browning in *Ctns*^−/−^ mice (as evidenced by the detection of inguinal WAT UCP1 protein and increased expression of beige adipose cell markers CD137, Tmem and Tbx1).^5,14^ Cox2 is a downstream effector of β-adrenergic signaling and induces biogenesis of beige cells in WAT depots.^32^ We showed that anakinra normalized inguinal WAT Cox2, Pgf2α as well as important inflammatory molecules (Tlr2, MyD88 and Trap6) expression in *Ctns*^−/−^ mice and normalized key inflammatory molecules (Tlr2 and MyD88) involved in adipose tissue browning in *Ctns*^−/−^ mice (**Figure 4**). Recent data suggest that IL1β signaling mediates adipocyte browning via regulating of mitochondrial oxidative responses in both cultured human and animal adipocytes.^33^

We studied the impact of anakinra on muscle wasting in *Ctns*^−/−^ mice. Anakinra normalized lean mass content, gastrocnemius wet weight as well as muscle function in *Ctns*^−/−^ mice relative to WT mice (**Figure 2, K to N**). Furthermore, anakinra normalized average cross-sectional area as well as muscle fat infiltration of gastrocnemius muscle in *Ctns*^−/−^ mice (**Figure 5, A-E**). Muscle fat infiltration is a significant predictor of both muscle function and mobility function across a wide variety of comorbid conditions such as diabetes, spinal cord injury and kidney disease.^34–36^ Muscle adipose tissue may release pro-inflammatory cytokines within the muscle and impair the local muscle environment, impair blood flow or increase the rate of lipolysis within skeletal muscle resulting in an increased concentration of glucose within the skeletal muscle itself followed by insulin resistance.^37,38^

We investigated the impact of anakinra on signaling molecules that modulate muscle mass regulation in *Ctns*^−/−^ mice. Systemic inflammation decreases muscle regeneration and increases muscle catabolism that eventually leads to muscle wasting in CKD. Inhibition of IL-1 reduced systemic inflammation (decreased serum CRP and IL-6) in CKD patients.^12,39^ A recent study also showed that blockade of IL-1 signaling significantly improved inflammatory status (a decrease in plasma concentration of IL-6 and TNF) and antioxidative properties in CKD patients.^40^ Systemic inflammation, as assessed by serum concentration of CRP, is a strong and independent risk factor for skeletal muscle wasting in CKD patients.^41^ IL-1 has been shown to stimulate the expression of catabolic genes.^10,42,43^ MAPK are a family of protein phosphorylating enzymes that regulate a diverse aspect of cellular responses including skeletal muscle regeneration and differentiation.^44–46^ Activation of the NF-κB signaling pathway leads to severe muscle wasting in mice.^47^ IL-1α and IL-1β, both isoforms of IL-1, stimulated catabolism in C2C12 myotubes via activation of NF-κB signaling and atrogin-1 expression.^7^ Importantly, anakinra attenuated or normalized phosphorylation of muscle ERK 1/2, JNK, p38 MAPK and NF-κB p65 protein content in *Ctns*^−/−^ mice. In addition, anakinra improved muscle regeneration and myogenesis by decreasing mRNA expression of negative regulators of skeletal muscle mass (Atrogin-1 and Myostatin) as well as increasing muscle mRNA expression of pro-myogenic factors (MyoD and Myogenin) in *Ctns*^−/−^ mice (**Figure 6**). Our findings were in agreement with a recent report demonstrating that IL-1β administration inhibiting muscle mRNA expression of atrogin-1.^9^

Finally, we studied the impact of anakinra on muscle transcriptomics in *Ctns*^−/−^ mice. We showed that anakinra normalized or attenuated muscle expression of 12 out of 20 top differentially expressed genes in *Ctns*^−/−^ mice. ^14^ Detailed functional significance of each of these 12 differentiated expressed muscle genes is listed in **Table 1**. Anakinra normalized (Csrp3, Ly6a, Nlrc3, Tnnc1 and Tpm3, Atf3, Cidea, Sncg and Tbc1d1) and attenuated (Cyfip2 and Fhl1) muscle gene expression in *Ctns*^−/−^ mice relative to WT mice **(Supplemental figure 1 & 2)**. Increased expression of Csrp3, Cyfip2, Fhl1, Nlcr3, Tnnc1 and Tpm3 have been implicated in muscle atrophy and muscle weakness.^48–53^ Increased expression of Ly6a has been associated with aberrant remodeling of the extracellular matrix during skeletal muscle regeneration.^54^ In addition, increased expression of Ankrd2 as well as decreased expression of Atf3 impairs muscle function and impairs survival of motor neuron and muscle innervation.^55–57^ Recent data also suggest that decreased expression of Cidea and Sncg are associated with higher energy expenditure as well as accelerated lipolysis of adipose tissue.^58,59^ Furthermore, decreased expression of Tbc1d1 impairs glucose transport in skeletal muscle and has been implicated in follistatin-induced muscle hypertrophy.^60^

## CONCLUSION

We report that anakinra attenuates adipose tissue browning and muscle wasting in *Ctns*^−/−^ via multiple cellular mechanisms (**Figure 7**). IL-1 receptor antagonist may represent a novel targeted treatment for cachexia in patients with INC by attenuating adipose tissue browning and muscle wasting.

**Figure 7:**
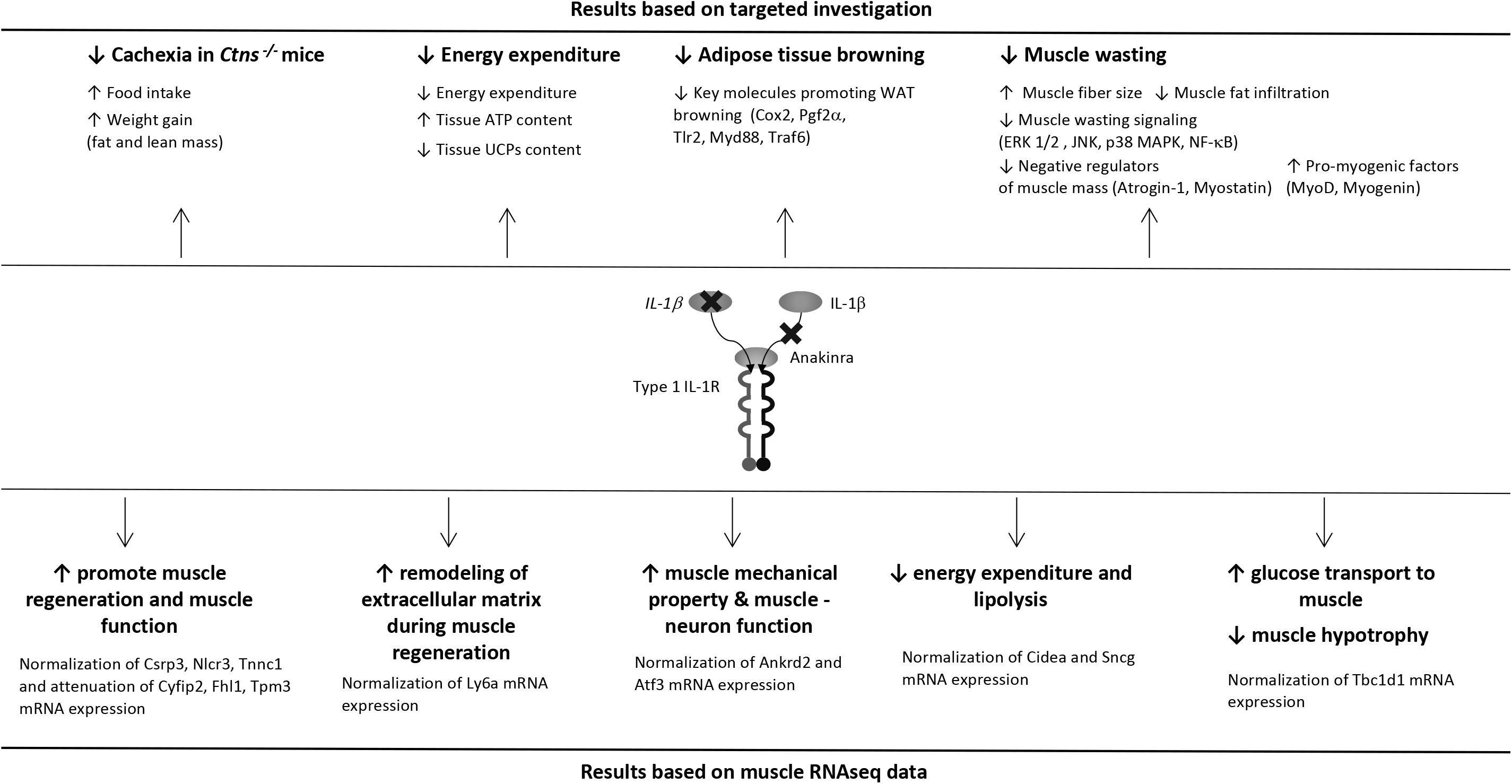
Summary of the beneficial effects of anakinra on cachexia, energy homeostasis, adipose tissue browning and muscle wasting in *Ctns*^−/−^ mice.

## ACKNOWLEDGMENTS

All authors of this manuscript certify that they complied with the ethical guidelines for authorship and publishing in the Journal of Cachexia, Sarcopenia and Muscle update 2017.^61^

## Supplemental information

**Supplemental Table 1:** Immunoassay information for blood and serum chemistry, muscle adenosine triphosphate content as well as muscle and adipose tissue protein analysis.

| <u>Blood & Serum chemistry</u> | <u>Assay information</u> |
| --- | --- |
| Bicarbonate & BUN | VetScan Comprehensive Diagnostic Profile, Abaxix, 500-0038 |
| Creatinine | LC-MS/MS method |
| <u>Muscle & adipose tissue</u> | <u>Assay information</u> |
| ATP Assay Kit (Colorimetric / Fluorometric) | Abcam, ab83355 |
| IL-1β | RayBiotech, ELM-IL1b-CL |
| IL-6 | RayBiotech, ELM-IL6 |
| Mouse, rat, human ERK1/2, JNK, p38 MAPK phosphorylation ELISA sampler kit | RayBiotech, CBEL-ERK-SK |
| Mouse NFkB p65 (phospho-Ser536) ELISA kit | RayBiotech, PEL-NFKBP65-S536-T-1 |
| Mouse total NFkB p65 ELISA kit | RayBiotech, PEL-NFKBP65-S536-T-1 |
| Mouse Ucp1 ELISA kits | Aviva Systems Biology, OKCD02970 |
| Mouse Ucp3 ELISA kits | Aviva Systems Biology, OKEH05259 |

**Supplemental Table 2S:**
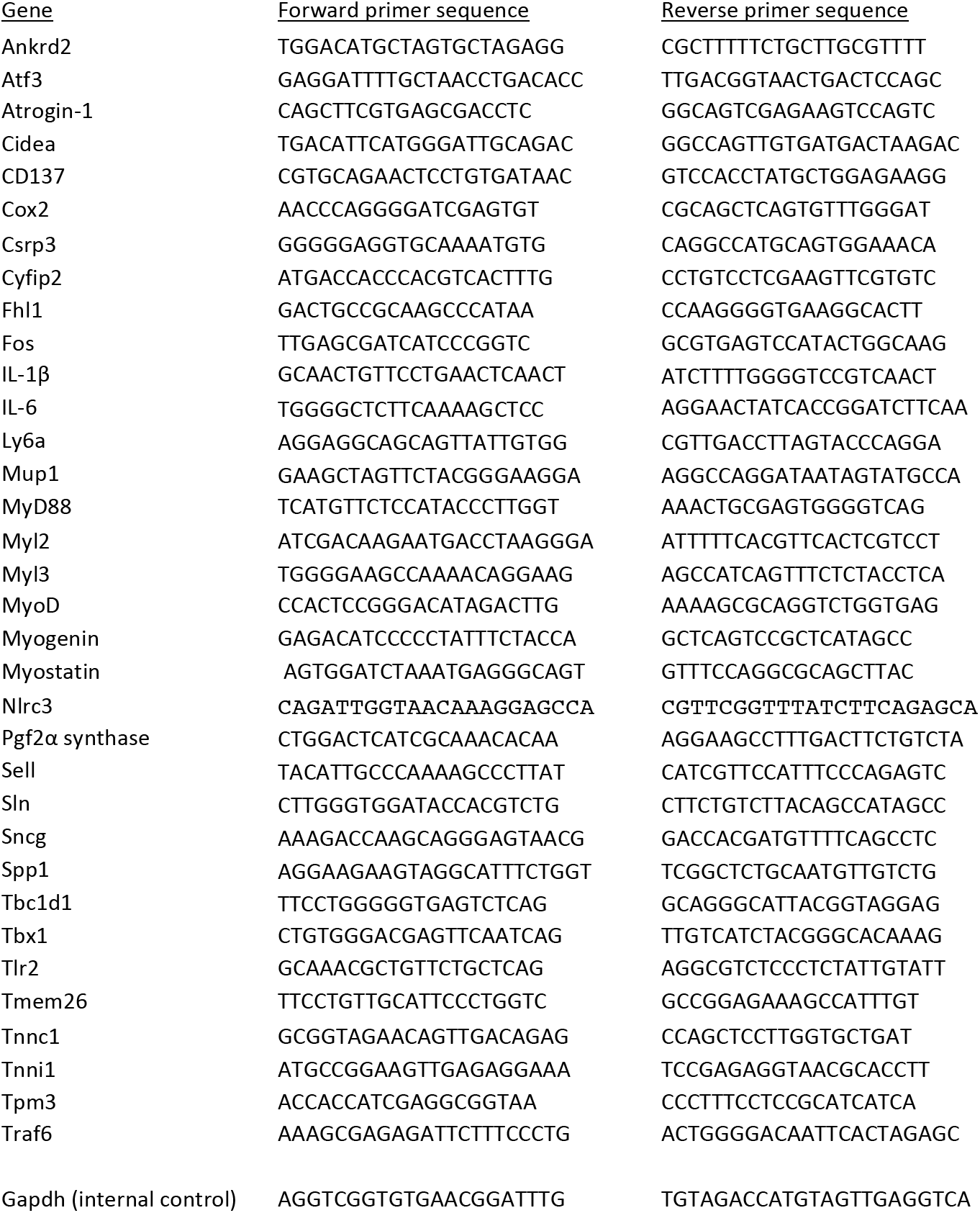
PCR primer information.

**Supplemental Table 3S:** Serum and blood chemistry of 12-month old *Ctns*^−/−^ and wild type control mice. Data are expressed as mean ± SEM. Result of serum chemistry of *Ctns*^−/−^ mice were compared to WT mice. ^*#*^ p<0.05, significantly increased in *Ctns*^−/−^ mice relative to WT mice.

|  | WT | *Ctns*<sup>-/-</sup> |
| --- | --- | --- |
|  | n = 12 | n = 12 |
| BUN (mg/dL) | 26.4 ± 3.5 | 56.7 ± 11.3 # |
| serum creatinine (mg/dL) | 0.06 ± 0.03 | 0.21 ± 0.05 # |
| Bicarbonate (mmol/L) | 27.3 ± 2.5 | 27.3 ± 3.2 |

**Supplemental Table 4S:** Serum and blood chemistry of *Ctns*^−/−^, *Ctns*^−/−^*Il6*^−/−^, *Ctns^−/−^Il1β*^−/−^ and wild type control mice. Experiment was 6 weeks. Data are expressed as mean ± SEM. Result of *Ctns*^−/−^, *Ctns*^−/−^*Il6*^−/−^ and *Ctns^−/−^Il1β*^−/−^ mice were compared to WT mice. ^*#*^ p<0.05, significantly increased in *Ctns*^−/−^, *Ctns*^−/−^*Il6*^−/−^ and *Ctns^−/−^Il1β*^−/−^ mice relative to WT mice.

|  | WT | *Ctns*<sup>-/-</sup> | *Ctns*<sup>-/-</sup>*Il6*<sup>-/-</sup> | *Ctns*<sup>-/-</sup>*Il1β*<sup>-/-</sup> |
| --- | --- | --- | --- | --- |
|  | n = 9 | n = 9 | n = 9 | n = 9 |
| BUN (mg/dL) | 23.6 ± 4.8 | 49.7 ± 4.3 # | 69.1 ± 7.7 # | 49.5 ± 3.5 # |
| serum creatinine (mg/dL) | 0.06 ± 0.01 | 0.17 ± 0.04 # | 0.20 ± 0.03 # | 0.14 ± 0.02 # |
| Bicarbonate (mmol/L) | 27.4 ± 3.8 | 26.9 ± 3.5 | 26.4 ± 3.2 | 27.3 ± 3.1 |

**Supplemental Table 5S:** Serum and blood chemistry of *Ctns*^−/−^ and wild type control mice. Twelve-month old WT and *Ctns*^−/−^ mice were treated with anakinra (2.5 mg.kg per day, IP) or vehicle (normal saline) for 6 weeks. Vehicle treated *Ctns*^−/−^ mice were fed *ad libitum* while other group of mice were pair-fed with the same amount of rodent diet as consumed by vehicle treated *Ctns*^−/−^ mice. Result of serum chemistry of *Ctns*^−/−^+Vehicle mice were compared to WT+Vehicle mice while results of *Ctns*^−/−^+Anakinra mice were compared to WT+Anakinra mice. Data are expressed as mean ± SEM. ^*#*^ p<0.05, significantly increased in *Ctns*^−/−^ + Vehicle and *Ctns*^−/−^ + Anakinra relative to WT + Vehicle and WT + Anakinra mice, respectively.

|  | WT + Vehicle | WT + Anakinra | *Ctns*<sup>-/-</sup> + Vehicle | *Ctns*<sup>-/-</sup> + Anakinra |
| --- | --- | --- | --- | --- |
|  | n = 9 | n = 9 | n = 9 | n = 9 |
| BUN (mg/dL) | 25.6 ± 3.1 | 24.7 ± 3.7 | 53.1 ± 13.2 # | 47.6 ± 7.2 # |
| serum creatinine (mg/dL) | 0.08 ± 0.02 | 0.09 ± 0.03 | 0.21 ± 0.05 # | 0.16 ± 0.03 # |
| Bicarbonate (mmol/L) | 26.7 ± 3.6 | 27.1 ± 2.7 | 27.3 ± 2.6 | 26.9 ± 3.7 |

**Supplemental figure 1:**
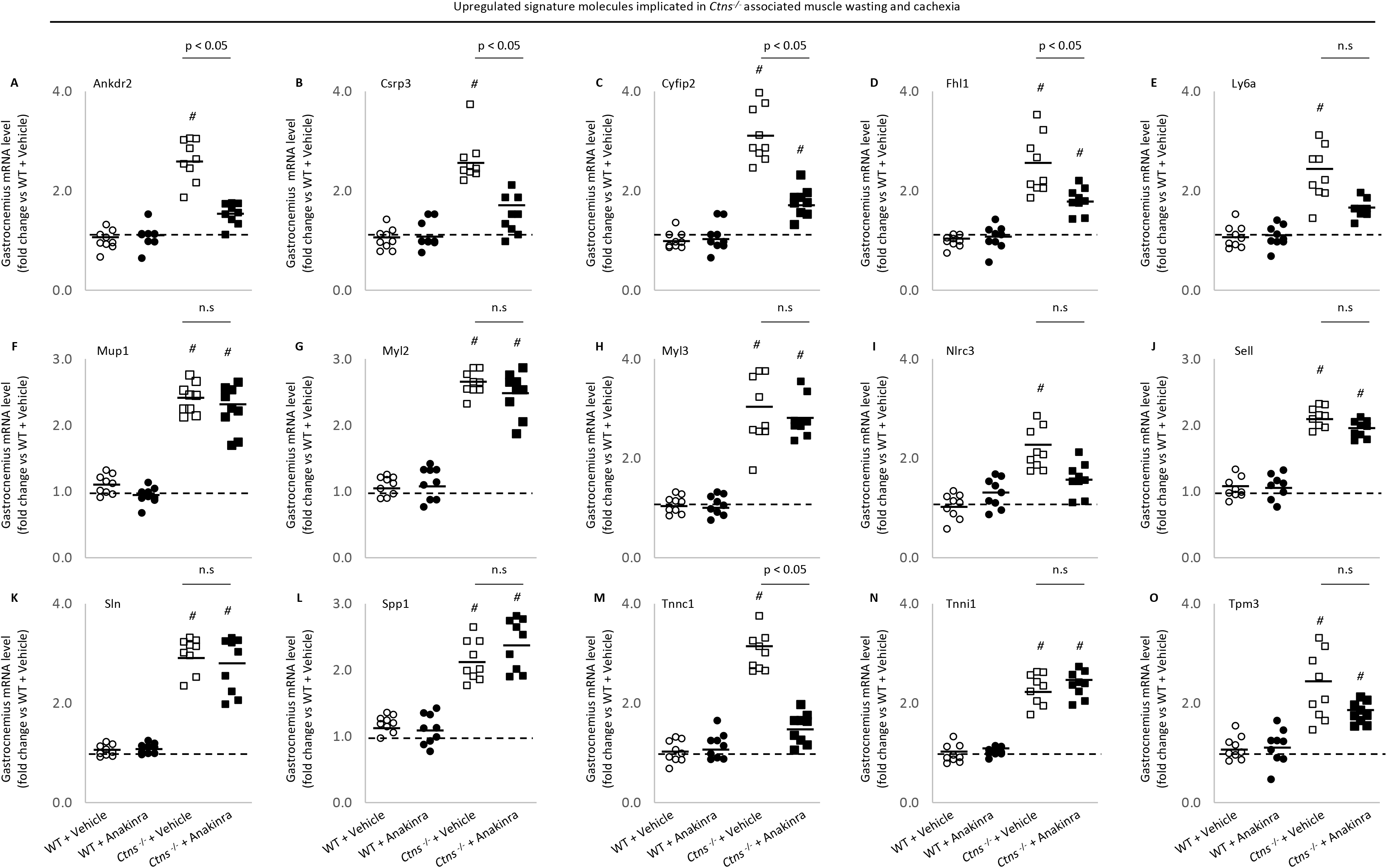
Anakinra attenuates upregulated signature molecules implicated in *Ctns*^−/−^ associated muscle wasting and cachexia. Gastrocnemius muscle expression of interested genes in mice was measured by qPCR. Final results were expressed in arbitrary units, with one unit being the mean level in vehicle-treated WT mice. Data are expressed as mean ± SEM. Results of vehicle-treated *Ctns*^−/−^ mice were compared to vehicle-treated WT mice while results of anakinra-treated *Ctns*^−/−^ mice were compared to that of anakinra-treated WT mice. In addition, results of anakinra-treated *Ctns*^−/−^ mice were compared to vehicle-treated *Ctns*^−/−^ mice. ^*#*^ p < 0.05.

**Supplemental figure 2:**
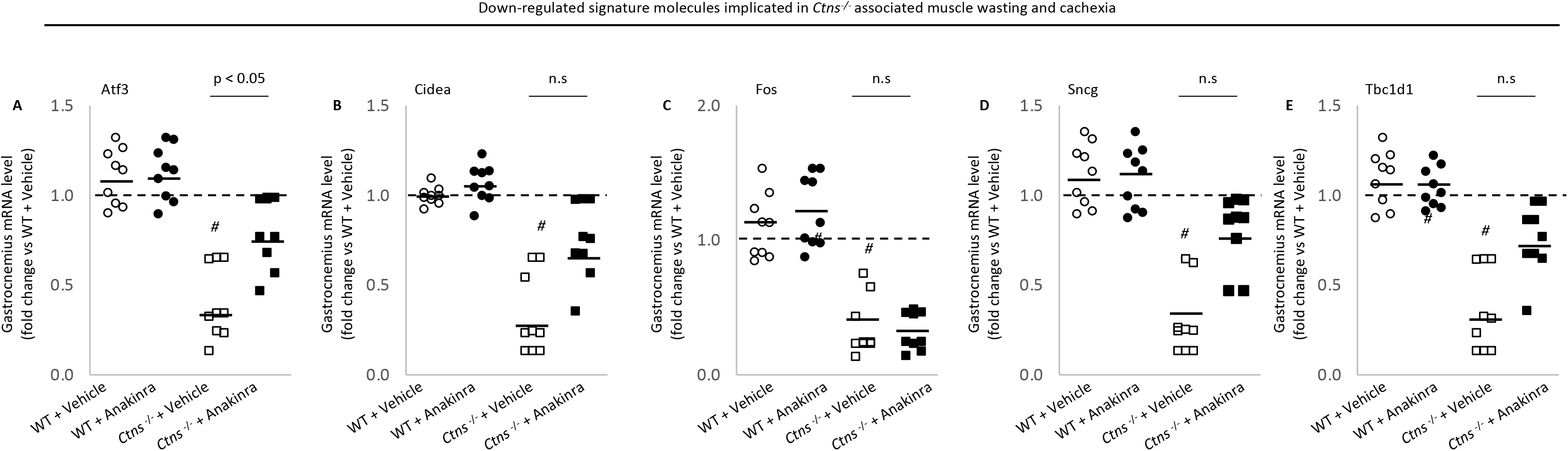
Anakinra attenuates downregulated signature molecules implicated in *Ctns*^−/−^ associated muscle wasting and cachexia. Gastrocnemius muscle expression of interested genes in mice was measured by qPCR. Final results were expressed in arbitrary units, with one unit being the mean level in vehicle-treated WT mice. Data are expressed and analyzed as supplemental figure 1.

